# Hematopoietic stem and progenitor cell heterogeneity is inherited from the embryonic hemogenic endothelium

**DOI:** 10.1101/2022.09.28.509963

**Authors:** Joey J. Ghersi, Gabriel Baldissera, Jared Hintzen, Stephanie A. Luff, Siyuan Cheng, Ivan Fan Xia, Christopher M. Sturgeon, Stefania Nicoli

## Abstract

Multipotent hematopoietic stem/progenitor cells (HSPCs) generate all mature blood cells in the erythroid, lymphoid, and myeloid lineages. HSPCs are initially produced in the embryo, via transdifferentiation of hemogenic endothelial cells (hemECs) in the aorta-gonad mesonephros (AGM). HSPCs in the AGM are functionally heterogenous in differentiation and proliferative output, but how these intrinsic differences are acquired remains unanswered. This knowledge could inform approaches to overcome the dysregulation of HSPC heterogeneity associated with poor outcomes of autologous transplants. Here we discovered that loss of microRNA (miR)-128 (miR-128^Δ/Δ^) in zebrafish leads to an expansion of hemECs forming replicative HSPCs in the AGM, and a skew towards the erythroid and lymphoid lineages in larval and adult stages. Furthermore, we found that inhibiting miR-128 during the differentiation of human pluripotent stem cells into hemECs, but not during the endothelial-to-hematopoietic transition, recapitulated the lineage skewing. *In vivo*, expression of wild-type miR-128 in endothelium restored the blood lineage distribution in miR-128^Δ/Δ^ zebrafish. We found that miR-128 represses the expression of the Wnt inhibitor *csnk1a1* and the Notch ligand *jag1b*, and thus promotes Wnt and Notch signaling in hemECs. De-repression of *cskn1a1* resulted in hemECs generating replicative and erythroid-biased HSPCs, whereas de-repression of *jag1b* resulted in hemECs forming lymphoid-biased HSPCs in the AGM and relative mature blood cells in adult. We propose that HSPC heterogeneity is established in hemogenic endothelium prior to transdifferentiation and is programmed in part by Wnt and Notch signaling modulation.

## Introduction

In the classical model of hematopoiesis, a homogeneous pool of hematopoietic stem cells (HSCs) proliferates while generating multipotent progenitors that, by following a stepwise restriction of lineage potential, generate all mature blood and immune cells (*1*). This model has been challenged by evidence for molecular and functional heterogeneity within the HSC pool. HSC transplantation, barcoding and fate mapping experiments showed that only a few HSCs can produce all blood cells, while the majority of HSC differentiation is restricted or imbalanced to few lineages (*2-7*). Furthermore, HSCs are different in their proliferative capacity which influence self-renewal kinetics (*8, 9*), with some HSCs been able to generate specific blood cells without undergoing cell division (*10*). Single-cell sequencing analysis confirmed that adult HSCs are an heterogenous mixture of multipotent stem and progenitor cells (HSPCs) having different cell cycle status, transcriptional lineage priming, and blood lineage outputs (*11, 12*). How HSPCs acquire these intrinsic phenotypic differences is currently unknown. This lack of knowledge is critical to understand how to regulate the production of HSPC *in vivo*, as well as *ex-vivo* where HSPC heterogeneity influences the success of autologous hematopoietic stem cell transplantation in clinic (*13, 14*).

Cell tracing experiments of arterial hematopoietic clusters during embryonic development showed that multiple HSPC clones develop and have long-term engraftment lineage biases (e.g lymphoid or myeloid) once they migrate in the definitive hematopoietic organs in juvenile and adult stages (*2, 5, 15*). HSPC heterogeneity is therefore observed in the embryo aorta-gonad mesonephros (AGM) (*16-18*) where nascent HSPCs (nHSPCs) are made via an endothelial-to-hematopoietic transition (EHT) from hemogenic endothelial cells (hemECs) (*19*). Even though hemECs are the direct precursor of nHSPCs it is unknown if they contribute to HSPC heterogeneity.

Here, using single cell RNA sequencing (scRNA-seq) and phenotypic analysis of AGM endothelial cells in nHSPC lineage priming models *in vivo* and *in vitro*, we discovered a previously unappreciated and unexpected mechanism in the endothelium that regulate HSPC heterogeneity prior to EHT.

## Results

### miR-128 regulates nascent HSPC heterogeneity

To search for vascular regulators of EHT, and potentially of HSPC heterogeneity, we screened for hematopoiesis defects in zebrafish embryos lacking the expression of miR-128, a highly conserved intronic miRNA enriched in endothelial cells (*20, 21*) and regulated in normal and malignant adult hematopoiesis (*22-25*). Notably, we observed an increased number of cells expressing the nHSPC marker *cmyb* in a mutant lacking expression of both *miR-128-1* and *miR-128-2* (hereafter, miR-128^Δ/Δ^) (*21*) (Fig.1 A and B). The expression of *r3hdm1* and *arpp21*, miR-128 host genes, was unchanged in miR-128^Δ/Δ^ (Supp Fig. 1A), suggesting direct contribution of miR-128 loss to the HSPC phenotype. Specifically, the number of HSPCs was first increased in miR-128^Δ/Δ^ relative to wild-type in the AGM during EHT at 32 hours post fertilization (hpf) and in the secondary hematopoietic organ, the caudal hematopoietic tissue (CHT), the equivalent of the fetal liver in mammals, at 3 days post fertilization (dpf) (Fig. 1A). Expansion of HSPCs was further noted at 6 dpf in the definitive hematopoietic organs, the thymus and the kidney marrow (KM), the equivalent of the bone marrow in mammals (Fig.1 B). Furthermore, *gata2b+* and *runx1+* hemECs, HSPC direct precursors in the AGM, were also increased in miR-128^Δ/Δ^ (Fig. 1C) while vascular development was normal (Supp Fig.1 B-G) supporting that HSPC expansion is linked to an increase in hemECs development and EHT.

**Figure. 1.**
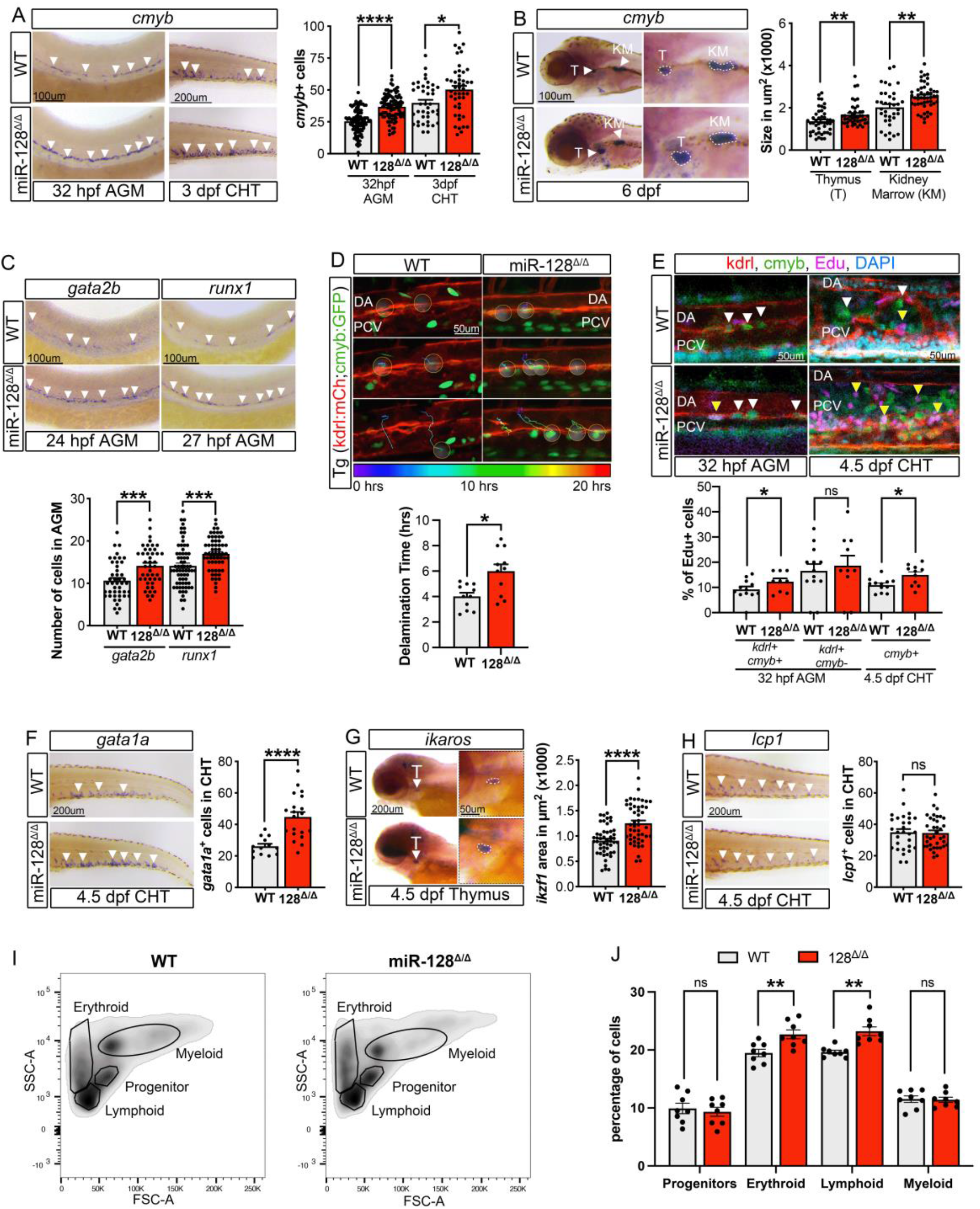
HSPC heterogeneity is altered in miR-128^Δ/Δ^. (A) Whole mount in situ-hybridization (WISH) against *cmyb* at 32 hpf and 3 dpf in wild-type (WT) or miR-128^Δ/Δ^ AGM and CHT. Quantification represent the number of cmyb^+^ cells in AGM or CHT. (B) WISH against *cmyb* at 6 dpf in WT and miR-128^Δ/Δ^ in thymus and kidney marrow. Quantification of the area of *cmyb*^+^ cells. (C) WISH against *gata2b* and *runx1* at 24 hpf and 27 hpf respectively of WT and miR-128^Δ/Δ^ AGM. Quantification of the number of *gata2b*^+^ and *runx1*^+^ cells. (D) Dragon-tail analysis of time-lapse imaging of WT and miR-128^Δ/Δ^ Tg(*kdrl:mCherry*^*s896*^,*cmyb:GFP*^*zf169*^) performed from 24 to 50 hpf. Circles represent newly nHSPCs (*kdrl*^+^,*cmyb*^+^). Quantification represents the delamination time in hours (hrs) of each traced cells. (E) Confocal images of Tg(*kdrl:mCherry*^*s896*^,*cmyb:GFP*^*zf169*^) wild-type and miR-128^Δ/Δ^ AGM at 32 hpf and CHT at 4.5 dpf after Edu incorporation. Replicative nHSPCs are increased (*kdrl*^+^,*cmyb*^+^, Edu^+^, yellow arrowheads and *kdrl*^+^,*cmyb*^+^, Edu^-^ white arrowheads) in miR-128^Δ/Δ^ while replicative endothelial cells are unchanged (*kdrl*^+^,*cmyb*^-^ Edu^+^). (F-H) WISH of (F) *gata1a*, (G) *ikaros* and (H) *lcp1* at 4.5 dpf with their quantification. Erythroid and lymphoid progenitors are increased in miR-128^Δ/Δ^. (I) Flow cytometry population analysis defined by SSC-A and FSC-A of 1-month-old dissected WT and miR-128^Δ/Δ^ whole kidney marrow. (J) Quantification of cell population identified by flow cytometry. Erythroid and Lymphoid fraction are increased in miR-128^Δ/Δ^ whereas progenitors and myeloid cells are unchanged, compared to WT. All Quantification are represented with mean ± SEM. ns: p>0.05, *p≤0.05, **p≤0.01, *** p≤0.001, **** p≤0.0001. Arrowheads indicate cells stained by WISH. Abbreviations: aorta gonad mesonephros (AGM), caudal hematopoietic tissue (CHT), thymus (T), kidney marrow (KM), dorsal aorta (DA), posterior cardinal vein (PCV), side scatter A (SSC-A), forward scatter A (FSC-A).

To assess the phenotypes of nHSPCs, we generated miR-128^Δ/Δ^ in the transgenic endothelial- and nHSPC-reporter line *Tg(kdrl:mCherry*^*s896*^*;cmyb:GFP*^*zf169*^) (18) and performed time-lapse live imaging throughout EHT, from 24 to 50 hpf. We found that compared to wild-type, miR-128^Δ/Δ^ *kdrl*+*cmyb*+ nHSPCs remained in the ventral floor of the dorsal aorta (DA) longer while dividing, thus showing a significant delay in delamination time (Fig.1 D and movie 1 A and B). To investigate potential differences in nHSPC proliferation, we injected embryos with 5-ethynyl-2’-deoxyuridine (Edu) to visualize cells undergoing DNA replication. We found that the number of proliferating nHSPCs (*kdrl*+*cmyb*+Edu+ cells), and the total number of nHSPCs (*kdrl+cmyb+* cells), were significantly increased in the AGM of miR-128^Δ/Δ^ versus wild-type at 32 hpf, whereas the number of proliferating endothelial cells (*kdrl*+Edu+ and *cmyb-* cells) was unchanged (Fig.1 E and Supp Fig.1 H). Similarly, the numbers of HSPCs (*cmyb*+ and *kdrl-*) and proliferating HSPCs (*cmyb*+ Edu+ and *kdrl-* cells) were increased in the 4.5 dpf CHT in miR-128^Δ/Δ^ versus wild-type (Fig.1 E, Supp Fig.1 I), supporting the increase in nHSPCs proliferation in miR-128^Δ/Δ^ embryos.

Next, we examined HSPC lineages in miR-128^Δ/Δ^. Critically, we found that erythroid and lymphoid progenitor cells were significantly expanded in 4.5 dpf miR-128^Δ/Δ^ CHT and thymus compared to wild-type (Fig.1 F and G), and their corresponding mature blood cells were also increased in 1-month-old whole kidney marrow (Fig.1 I and J). In contrast, the myeloid progenitors and lineages were unchanged in miR-128^Δ/Δ^ compared to wildtype (Fig.1 H-J).

Overall, our data indicate that loss of miR-128 led to an expansion of hemECs transdifferentiating in replicating nHSPCs, an expansion of erythroid and lymphoid progenitor cells in secondary hematopoietic organs, and altered blood composition in adulthood.

### Heterogeneous nHSPC populations are generated in the AGM and regulated by miR-128

To characterize EHT progression on a molecular level, we performed scRNA-seq of 22,230 *kdrl*+ endothelial cells isolated from the trunk of wild-type and miR-128^Δ/Δ^ 26 hpf embryos undergoing EHT (Supp Fig.2 A and B, and Supp table 1A). Of these cells, we focused on 6096 cells that express known vascular, arterial, and hematopoietic markers, composed of nine different clusters (C) that represent the continuous progression of EHT in both wild-type and mutant cells (Supp. Fig. 2 C-F and Supp table 1B and C). We identified tip cells (C0), and two arterial clusters (C1 and C2). C1 compared to C2 had an higher percentage of putative hemogenic endothelial cells expressing *gata2b* and *runx*1, thus was defined as a pre-hemogenic cluster (Fig. 2 A and B, Supp Fig.2 G and H, Supp table 1C and D). Accordingly, cells adjacent to C1 (C4) was composed mostly of cells expressing a continuum of hemogenic (*gata2b*/*runx1*) and nHSPCs (*cmyb*) markers, and therefore were defined as hemogenic endothelial cells undergoing the EHT (Fig.2 A and B, Supp Fig.2 G and H).

**Figure. 2.**
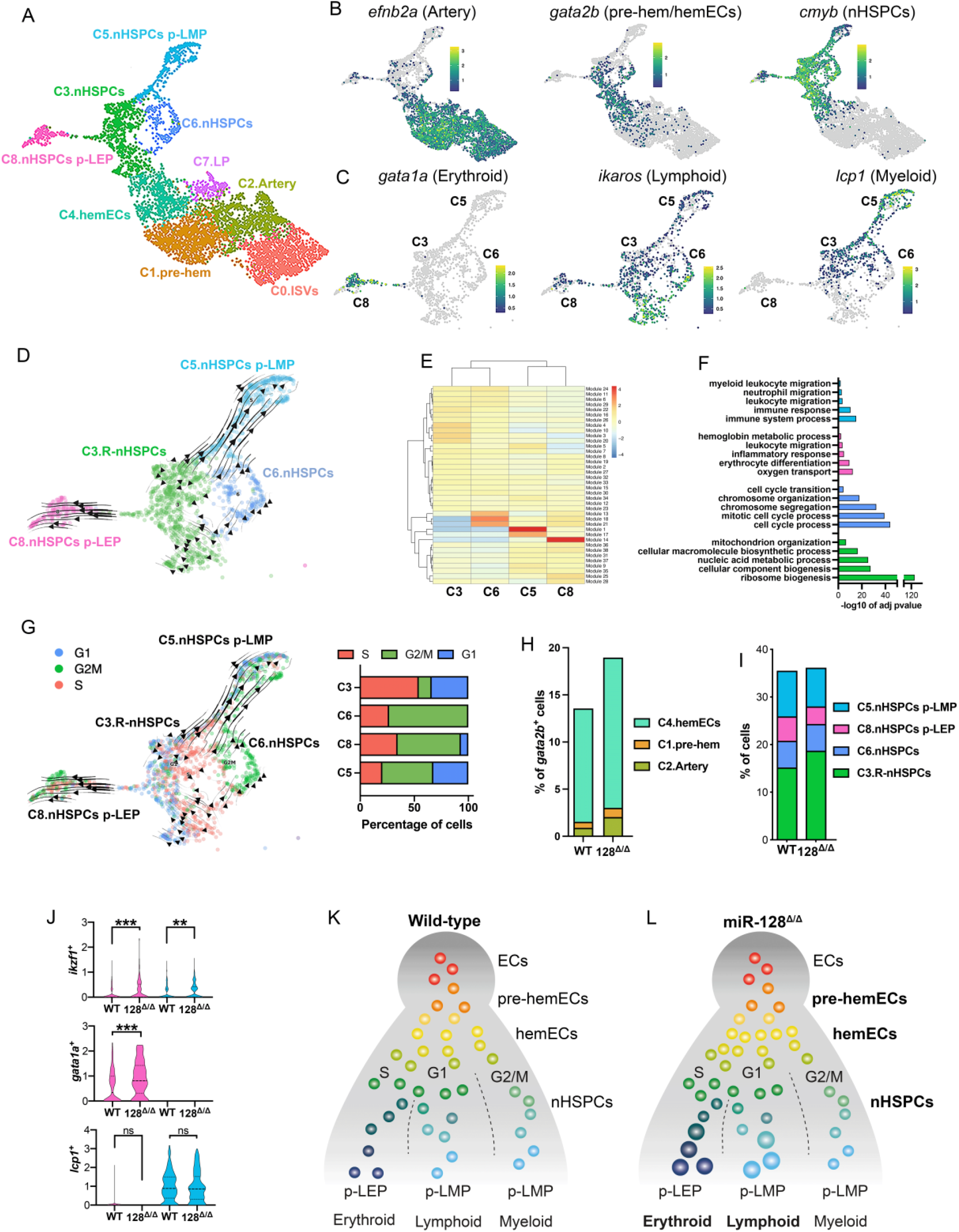
nHSPC heterogeneity is defined by cell cycle and lineage bias during embryonic EHT. (A) UMAP of defined EHT cluster cells from *kdrl+* trunk WT and miR-128^Δ/Δ^ ECs at 26 hpf. (B) UMAP representing *efnb2a, gata2b* and *cmyb* expression normalized with Z-score. (C) UMAP of *gata1a, ikaros* and *lcp1* expression normalized with Z-score, in nHSPC *cmyb*^+^ clusters. (D) RNA velocity trajectories in nHSPC clusters showing C3.R-nHSPCs progressing in LEP and LMP primed nHSPCs, while C6.nHSPCs progression is restricted to LMP primed nHSPCs. (E) Module analysis representing the differentiated set of defining genes for C3, C6, C5 and C8. (F) GO term analysis of module 1, 10, 14 and 18 revealed the gene expression signature differences between nHSPCs C3, C6, C5 and C8. (G) Cell cycle analysis on nHSPC clusters. Quantification of S, G2/M and G1 phase in C3, C6, C8 and C5. C3-nHSPCs cells are mainly in S phase and are indicated as replicative (R) and G1, while C6-nHSPCs in G2/M. (H) Percentage of *gata2b*+ cells in pre-EHT clusters (C2, C1 and C4) showing an increase of *gata2b+* cells. (I) Percentage of cells in nHSPCs clusters per genotype, showing an expansion of C3-R-nHSPCs in miR-128^Δ/Δ^ compared to WT cells. (J) Violin plot of *ikzf1, gata1a* and *lcp1* expression in clusters C8 (pink) and C5 (light blue) nHSPCs per genotype. (K) Model of nHSPC heterogeneity acquired during EHT in the AGM of WT embryos. (L) Model of nHSPC heterogeneity in miR-128^Δ/Δ^ showing nHSPCs heterogeneity skewed toward C3.R-nHSPCs, leading to the increase erythroid and lymphoid progenitors. ns: p>0.05, **p≤0.01, *** p≤0.001. Abbreviations: primed lympho-erythroid progenitors (p-LEP), primed lympho-myeloid progenitors (p-LMP), lymphatic progenitor (LP), Inter-segmental vessels (ISVs).

Cells emerging from the EHT C4 were grouped in four different nHSPC clusters that have higher expression of *cmyb* and lower expression of the vascular marker *kdrl* (Fig.2 A and B and Supp Fig.2 G and H). Two of these clusters showed high co-expression of genes priming lymphoid-erythroid progenitors (C8-nHSPC primed LEP) or lymphoid-myeloid progenitors (C5-nHSPC primed LMP) (Fig.2 A and C).

Importantly, these C8- and C5-lineage primed nHSPC branched from two *cmyb*+ clusters that differed in the expression of cell cycle genes, which we named C3- and C6-nHSPCs (Fig.2 D-F and Supp table 1E). Accordingly, when we ordered cells in each cluster according to the expression of cell cycle phase progression genes, we found that C3-nHSPC contained cells mainly in S phase whereas C6-nHSPC contained cells mainly in G2/M phase (Fig.2 G and Supp table 1F).

These data led us to hypothesize that nHSPCs arise from the EHT as a continuum of cells with different proliferation outputs and priming for distinct blood lineages. To test this, we performed an RNA-velocity analysis, which predicts the future state of individual cells by comparing the levels of unspliced and spliced mRNAs in scRNA-seq data (*26*). We used this approach to predict the future state of C3- and C6-nHSPCs. The data suggested that C3-nHSPCs, which we defined as replicative (R-nHSPCs), progresses into C8-nHSPCs primed LEP and C5-nHSPCs primed LMP (Fig.2 D and G) but the C6-nHSPCs progresses only into C5-nHSPCs primed LMP (Fig. 2 D and G). Our data suggest that within the AGM nHSPCs are born with distinct cell cycle states that enable their priming in different hematopoietic lineages (Fig.2 K).

We then tested how miR-128 loss influences nHSPCs heterogenicity in the AGM. Relative to wild-type, miR-128^Δ/Δ^ *kdrl*+ cells had an expanded population of *gata2b+* cells in pre-EHT clusters (C2, C1 and C4), an increase of R-nHSPCs (C3) and an elevated expression of lymphoid (*ikzf1*) and erythroid (*gata1a*) markers in C8-nHSPCs primed LEP and C5-nHSPCs primed LMP, whereases myeloid marker (*lcp1*) was unchanged (Fig.2 H-J and Supp table G). Thus, miR-128 limits hemECs generating of R-nHSPCs and their subsequent lymphoid and erythroid lineage priming (Fig.2 L).

### miR-128 inhibits HSPC heterogeneity with hemECs but not HSPCs

To investigate the expression of miR-128 during development, we performed fluorescent activated cell sorting of wild-type endothelial, non-endothelial cells and nascent HSPCs using *Tg(kdrl:mCherry*^*s896*^*;cmyb:GFP*^*zf169*^) embryos at 26 hpf followed by RT-qPCR. Intriguingly, we found that miR-128 expression was higher in endothelial cells (*kdrl*+*cmyb-)* than in nHSPCs (*kdrl*+*cmyb*+) and non-endothelial (*kdrl*-*cmyb-)* cells (Supp Fig.3 A). Moreover, the expression of *r3hdm1* and *arpp21*, the genes hosting miR-128, were found to be enriched in pre-EHT endothelial clusters in our scRNA-seq data (Supp Fig.3 B). Correspondingly, introduction of a wild-type copy of miR-128 under the control of the vascular-promoter *fli1a (27)* into miR-128^Δ/Δ^ rescued the bias of the blood lineages in the CHT (Fig.3 A-C, Supp Fig.3 C). Based on these data, we hypothesized that the expansion of nHSPC heterogenity in miR-128^Δ/Δ^ AGM is due to an earlier loss of miR-128 expression in hemEC precursors.

**Figure. 3.**
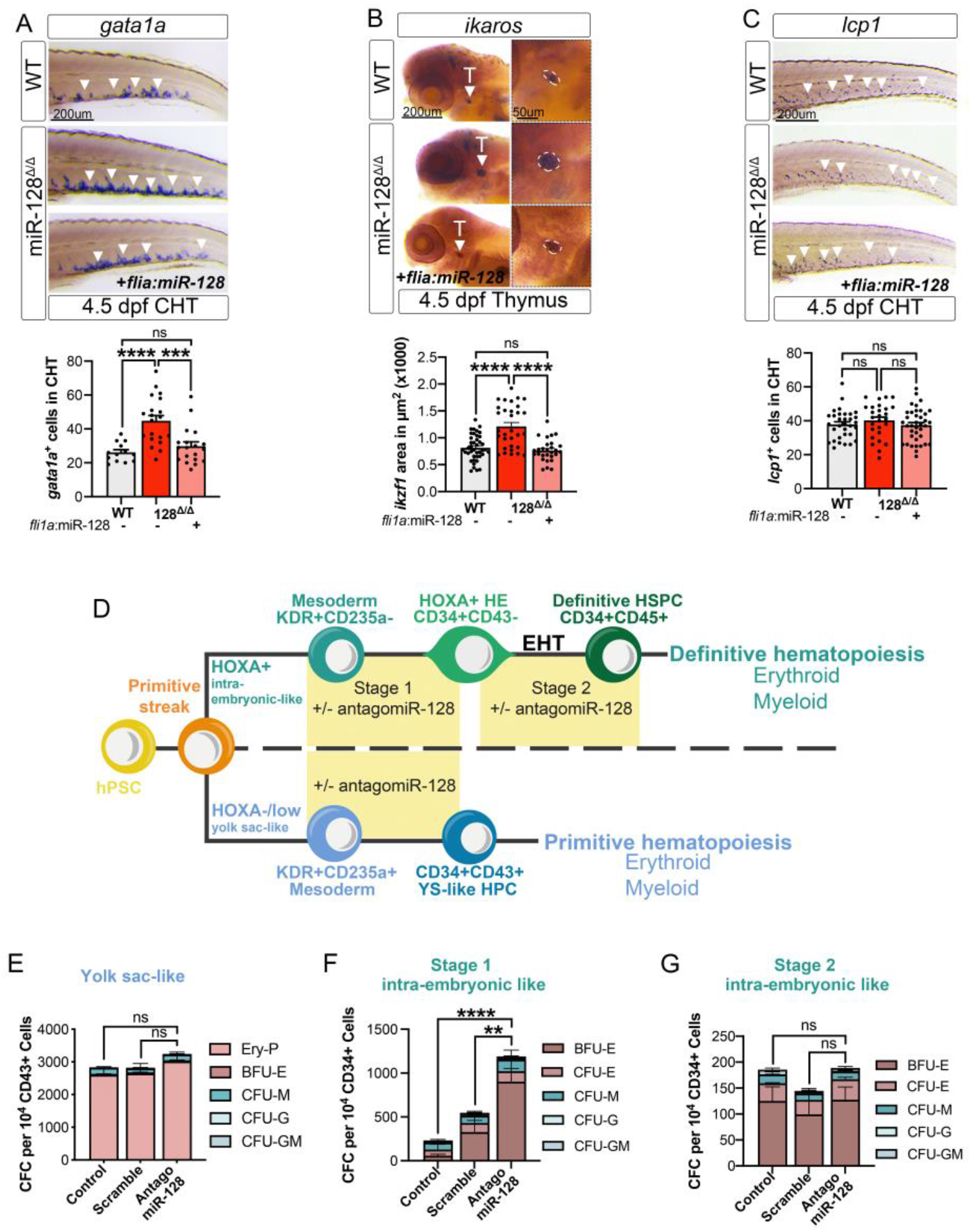
Endothelial miR-128 regulates HSPC heterogeneity. (A-C) WISH of (A) *gata1a*, (B) *ikaros* and (C) *lcp1* at 4.5 dpf and relative cells quantification as indicated. Quantification of *gata1a* in WT and miR-128^Δ/Δ^ are the same as in figure 1D. *fli1a*-endothelial expression of *miR-128* wild-type gene rescues to WT level the increase of erythroid and lymphoid progenitors of miR-128^Δ/Δ^. (D) Schematic of HSPC development *in vitro* using human pluripotent stem cells (hPSC) and treated as indicated with antagomiR-128 or scramble miR as control at different cell stages. (E) Colony forming cell quantification of erythroid (Ery-P, BFU-E) and myeloid (CFU-M, CFU-G, CFU-GM) of *HOXA/low-* program (primitive hematopoiesis). (F-G) Colony forming cell assay quantification of erythroid (BFU-E, CFU-E) and myeloid (CFU-M, CFU-G, CFU-GM) of *HOXA+* program (definitive hematopoiesis) during stage 1 (F) or stage 2 (G). All Quantification are represented with mean ± SEM. ns: p>0.05, **p≤0.01, **** p≤0.0001. Abbreviations: human pluripotent stem cells (hPSC), endothelial to hematopoietic transition (EHT), hematopoietic stem and progenitor cells (HSPC), hemogenic endothelium (HE), colony forming cell (CFC), caudal hematopoietic tissue (CHT), burst forming units erythroid (BFU-E) and colony forming units (CFU) of erythroid (E), granulocyte (G), myeloid (M), and mixed granulocyte/myeloid (GM).

To test this hypothesis, we employed an *in vitro* system using human pluripotent stem cell (hPSC) differentiation to recapitulate the earliest stages of hematopoietic development via EHT (Fig.3 D). Briefly, we have previously demonstrated that treating primitive streak-like cells (orange) with a CHIR99021 specifies a KDR+CD235a-mesodermal population, which in turn gives rise to a *HOXA+*CD34+CD43-population which harbors intra-embryonic-like hemECs (stage 1) (*28, 29*). These cells, in turn, undergo the EHT to form CD34+CD45+ definitive HSPCs (stage 2) (*28, 30-32*). Although these HSPCs lack engraftment potential in a xenograft model, a clonal multi-lineage assay (*32*) demonstrated that 10% of these hemECs possesses *bona fide* erythro-myelo-lymphoid multi-lineage potential (*32-46*). Thus, these HSPCs can be assessed for their ability to generate definitive erythroblasts and myeloid cells (Fig.3 D light green) (*32, 47, 48*).

In contrast, treating primitive streak-like cells with IWP2 and ACTIVIN A specifies a KDR+CD235a+ mesodermal population, which in turn gives rise to CD34+CD43+ *HOXA-* yolk sac-like hematopoietic progenitor cells, which can be assessed for their ability to give rise to primitive erythroblasts and myeloid cells (Fig.3 D light blue) (*31, 49-51*).

Using this hPSC model, we used an antagomir (*52*) to reduce miR-128 expression at different stages of *in vitro* hematopoietic development (Supp Fig.3 D). hPSCs treated with or without antagomir (or scramble miR control) during the differentiation of CD235a+ mesoderm into yolk sac-like hematopoietic progenitor cells showed no changes in overall hematopoietic output, and the cultures retained a similar lineage bias (Fig.3 D and E and Supp Fig.3 E-G), suggesting that miR-128 does not play a role in the emergence of the primitive hematopoietic program. In striking contrast, antagomir treatment during stage 1 of differentiation, during the specification of *HOXA+* hemECs from mesoderm, resulted in those cells exhibiting an overall 2-5 fold increase in definitive erythroid, but not myeloid, output (Fig.3 F and Supp Fig.3 H-J). Notably, inhibition of miR-128 after *HOXA+* hemEC specification, during the EHT in stage 2 of differentiation, did not affect the resultant output of CD34+CD45+ cells, nor their lineage bias (Fig.3 G and Supp Fig.3 K-L). Overall, these data suggest that miR-128 functions in endothelial cells, prior to EHT, to limit the generation of lineage biased HSPCs.

### HSPC heterogeneity can be modulated by miR-128-dependent Wnt and Notch endothelial activity prior EHT

To discern how miR-128 regulates HSPC diversity, we first compared the transcriptomes of 26 hpf *kdrl*+ cells from trunk miR-128^Δ/Δ^ and wild-type embryos, which include hemECs in the AGM (Supp Fig.4 A). We also used *kdrl*+ cells from heads as control of specificity. Multiple signaling components were de-regulated specifically in miR-128^Δ/Δ^ trunk *kdrl*+ cells (Supp Fig.4 A-C and Supp table 2A-C). As proof-of-principle we focused on miR-128-regulation of Notch and canonical Wnt signaling, key pathways of *in vivo* and *ex-vivo* HSPC production, predominantly active prior EHT (Supp Fig.4 D). We identified the Wnt regulator *csnk1a1* (*53*) and Notch regulator *jag1b* (*54-56*) as candidate targets of miR-128 prior EHT. These genes were highly expressed in arterial and pre-hemEC clusters (C2 and C1) (Supp Fig.4 E and Supp table 3A) and were responsive to changes in miR-128 activity. *csnk1a1*and *jag1b* were de-repressed in trunk miR-128^Δ/Δ^ cells and repressed by ectopic miR-128 endothelial expression in wild-type (Supp Fig.4 F). Moreover, their levels were normalized to wild-type when miR-128 endothelial expression was reintroduced in miR-128^Δ/Δ^ (Supp Fig.4 F), suggesting that *csnk1a1*and *jag1b* regulation in endothelia cells is miR-128 dependent.

To disrupt miR-128-mediated regulation of these targets, we used CRISPR/Cas9 to introduce indels into the miR-128 binding sites of the *csnk1a1* or jag1b genomic (g) 3’UTRs (Supp Fig.4 G). As predicted, the genetic perturbations led to de-repression of the associated transcripts at 24 hpf *in vivo* (Supp Fig.4 G). To determine if miR-128 regulation of *csnk1a1* or *jag1b* impacted the Wnt and Notch signaling pathways, we assessed the *Tg(TCF:nls-mCherry)*^*ia5*^ Wnt and *Tg(TP1:eGFP)*^*um14*^ Notch reporter lines, respectively. Notably, *csnk1a1* g3’UTR mutants showed an increase in Wnt-negative cells, *jag1b* g3’UTR mutants showed an increase in Notch-low cells in the ventral floor of the DA (the AGM), and miR-128^Δ/Δ^ presented both these phenotypes (Fig.4 A and B). In contrast, *kdrl*+ cells in the dorsal floor of the dorsal aorta, where the EHT is absent, had similar Wnt and Notch activity in all the genotypes (Supp Fig.4 H and I). Additionally, conserved Notch and Wnt-signaling targets had reduced expression in pre-EHT cluster cells (C2, C1 and C4) in miR-128^Δ/Δ^ and *csnk1a1* or *jag1b* g3’UTR mutants (Supp Fig. 4J and K, Supp table 3B). These targets were conserved across species, as these genes were also diminished in aorta-like CD34+ cells derived from the hPSCs treated with the miR-128 antagomir during stage 1 (Supp Fig.4 L and Supp table 4). Overall, these data suggest that direct post-transcriptional repression of *csnk1a1* and *jag1b* by miR-128 contributes to the inhibition of Wnt and Notch signaling prior EHT.

**Figure. 4.**
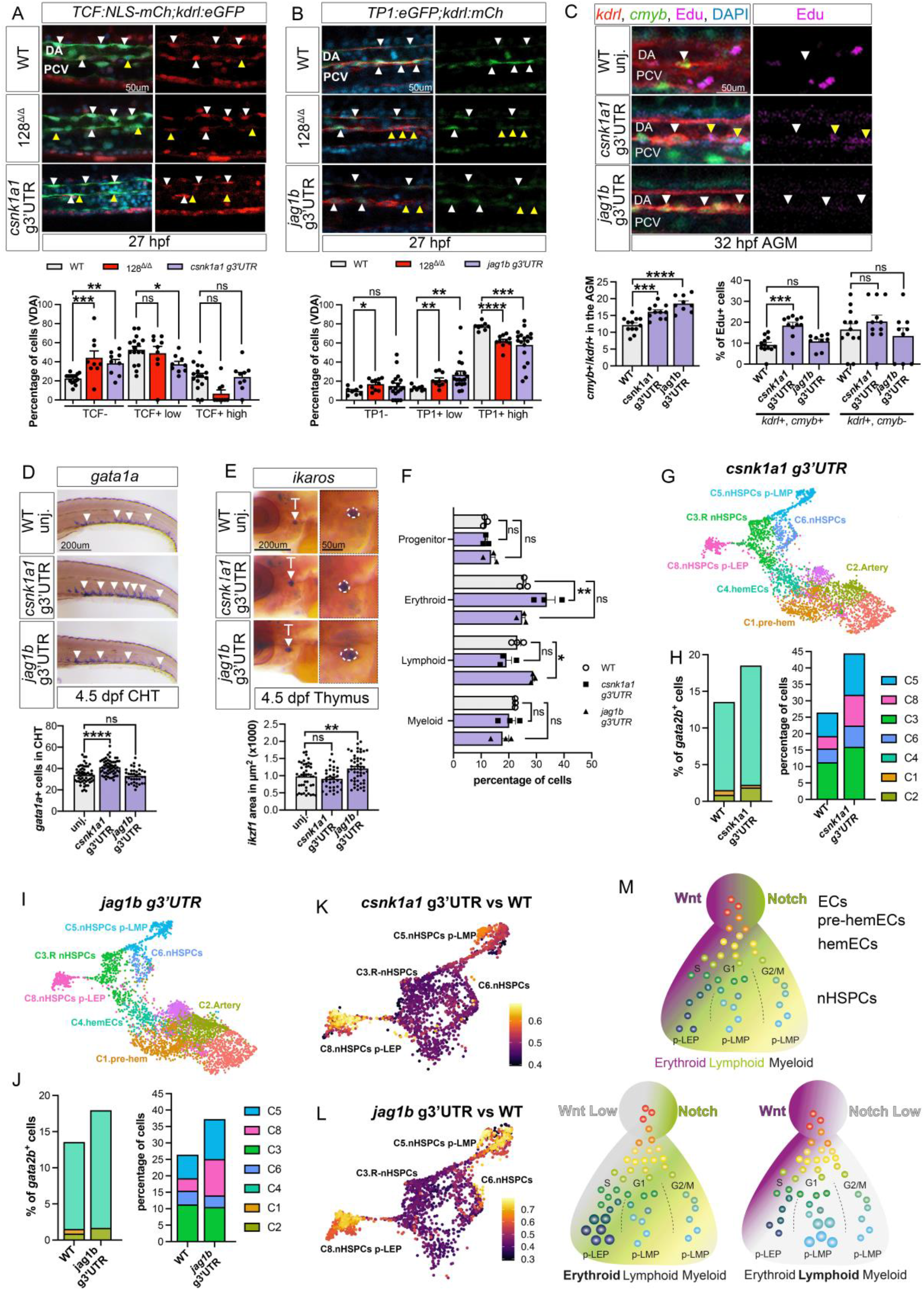
Endothelial jag1b-Notch and csnk1a1-Wnt signaling differentially regulate HSPC heterogeneity in the AGM. (A-B) Confocal lateral view of trunk 27 hpf of (A) Wnt (*TCF:NLS-mCherry*^*ia5*^,*kdrl:eGFP*^*zn1*^) and (B) Notch (*TP1:eGFP*^*um14*^,*kdrl:mCherry*^*s896*^) signaling through immunofluorescence. Quantification of Wnt and Notch classified as negative, low, and high cells based on the intensity of mCherry and GFP staining respectively. Arrowheads represent *kdrl*+, TCF+ or TP1+ high (white) or low/negative (yellow) cells. Cell quantification is reported in the ventral floor of the dorsal aorta (VDA). (C) Confocal images of Tg(*kdrl:mCherry*^*s896*^,*cmyb:GFP*^*zf169*^) wild-type and g3’UTR mutants AGM at 32 hpf. Replicative nHSPCs are increased (*kdrl*^+^,*cmyb*^+^ Edu+, yellow arrowheads) while replicative ECs (*kdrl*^+^,*cmyb*^-^, Edu+) are unchanged in the *csnk1a1* g3’UTR. Replicative nHSPCs (*kdrl*^+^,*cmyb*^+^, Edu+) and ECs (*kdrl*^+^,*cmyb*^-^, Edu+) are unchanged in the *jag1b* g3’UTR (*kdrl*^+^,*cmyb*^-^ Edu+). Total nHSPCs (*kdrl*^+^,*cmyb*^+^, white arrowheads) were increased in both genotype. (D,E) Quantification of the (D) erythroid and (E) lymphoid progenitors by WISH against *gata1a* and *ikaros* respectively, at 4.5dpf CHT or thymus of WT, *csnk1a1* g3’UTR or *jag1b* g3’UTR. (F) Quantification of adult blood cell population identified by flow cytometry of 2 month-old KM dissected from WT, *csnk1a1* g3’UTR or *jag1b* g3’UTR. Erythroid fraction is increased in *csnk1a1* g3’UTR and lymphoid fraction are increased in *jag1b* g3’UTR whereas progenitors and myeloid cells are unchanged, compared to WT. (G) UMAP of *csnk1a1* g3’UTR single *kdrl*+ trunk endothelial cells clustered based on defined EHT gene markers. (H) Percentage of *gata2b*+ cells and total cells in nHSPC clusters per genotype. (I) UMAP of *jag1b* g3’UTR single *kdrl*+ trunk endothelial cells clustered based on defined EHT gene markers. (J) Percentage of *gata2b*+ cells and total cells in nHSPCs cluster per genotype. (K and L) Analysis of experimental perturbation among genotype using MELD (see methods) assessed by the comparison between, *csnk1a1* g3’UTR versus WT (K) and *jag1b* g3’UTR versus WT (L). (M) Proposed model of HSPCs heterogeneity regulated by vascular hemogenic cells prior endothelial to hematopoietic transition. During pre-hemogenic or hemogenic development vascular signaling, like Wnt (dark purple) and Notch (green) limit the production of an heterogenous pool of HSPCs in the AGM. Diminishment of Wnt (light grey), like after de-repression of *csnk1a1*, results in an increase of replicative, and erythroid biased HSPCs in the AGM and relative erythroid cells in adult. Diminishment of Notch (light grey), like after de-repression of *jag1b*, increases lymphoid biased HSPCs in the AGM and relative lymphoid cells in adult stage. All Quantification are represented with mean ± SEM. ns: p>0.05, *p≤0.05, **p≤0.01, *** p≤0.001, **** p≤0.0001. Abbreviations: nuclear localization site (NLS), mCherry (mCh), aorta gonad mesonephrons (AGM), caudal hematopoietic tissue (CHT), primed lympho-erythroid progenitors (p-LEP), primed lympho-myeloid progenitors (p-LMP).

To discern the specific role of miR-128-mediated Wnt and Notch regulation in nHSPC heterogeneity we analyzed nHSPC phenotypes identified in miR-128^Δ/Δ^. Notably, *csnk1a1* g3’UTR mutants showed an expansion of replicative HSPCs (*kdrl+cmyb+*Edu+) in the AGM and only erythroid progenitors in the CHT at 4.5 dpf and KM at 2 months (Fig.4 C-F), whereas the *jag1b* g3’UTR mutant showed an expansion of HSPCs (*kdrl+cmyb+*Edu-) and lymphoid progenitors in the thymus at 4.5 dpf and KM at 2 months (Fig.4 C-F), supporting that different vascular signaling pathways control different HSPC phenotypes and relative blood cells. To verify HSPC heterogeneity in each g3’UTR mutants AGM, we performed scRNA-seq of *kdrl+* trunk cells at 26 hpf (Supp Fig.4 M-O and Supp table 5). The *csnk1a1* g3’UTR mutants showed an expansion of *gata2b+* cells in pre-EHT clusters, of C3 replicative nHSPCs, as well as of the C8 nHSPCs primed LEP and C5 nHSPCs primed LMP populations (Fig. 4G and H). Whereas *jag1b* g3’UTR mutants showed an expansion of *gata2b+* cells and nHSPCs primed LEP and LMP only (Fig.4 I and J and Supp table 5B and C). Interestingly, differential abundance testing on single nHSPCs (*57*) showed that de-repression of *csnk1a* had the largest effect on nHSPCs LEP primed (C8) whereas de-repression of *jag1b* affected both LEP and LMP primed nHSPCs (C8 and C5) (Fig.4 K and L). This is consistent with the increase in erythroid and lymphoid progenitor cells observed in the CHT of *csnk1a1* g3’UTR or *jag1b* g3’UTR mutants, respectively (Fig.4 D and E).

Overall, these results suggest that HSPCs heterogeneity is differentially established by multiple endothelial pathways, like Wnt and Notch, prior to EHT (Fig.4 M).

## Discussion

In this study, we discovered the regulatory networks in the vasculature that govern HSPC heterogeneity. This finding suggests that HSPCs inherit distinct behaviors, like cell cycle states and lineage priming, from AGM hemogenic endothelium that influence both embryo and adult blood composition. We found that different HSPC primed phenotypes originate from the activities of distinct signaling pathways in hemogenic endothelial cells prior to the EHT. Mechanistically, we showed that miR-128 post-transcriptional regulation of the canonical Wnt inhibitor *csnk1a1* limits the formation of replicative nHSPCs, erythroid-biased nHSPCs and progenitors. Correlatively, the miR-128 repression of the Notch ligand *jag1b* limits the generation of lymphoid-biased nHSPCs and progenitor cells in embryo and adult stage. Thus, Wnt and Notch signaling are differentially required to direct HSPCs proliferative and differentiation outputs influencing adult blood composition (Fig.4 M).

Heterogeneity in HSPCs has been observed in adult bone marrow as well as in embryonic hematopoietic organs, including the AGM. Interestingly, the regulation of HSPC heterogeneity is strongly associated with multisystem disease susceptibility and acquired genetic mosaicism during aging (*11, 58*). Whether HSPC heterogeneity can be “corrected” to improve these disease outcomes is yet to be considered since, until now, it was unknown how intrinsic HSPC heterogeneity can be regulated. Our discovery fills this gap of knowledge. Since AGM HSPCs are destined to generate blood over a lifetime, our discovery suggests that specific endothelial cell signaling pathways can be manipulated to either rebalance blood and immune cells or increase the production of one blood lineage vs another at birth. Either way, such modulation *in vivo* and *in vitro* might open new intervention avenues to modulate blood as needed, without compromising vascular niche dependent phenotypes.

Furthermore, HSPCs phenotypes in the AGM can functionally influence the blood composition of young adult animals. For example, lymphoid lineages are generated from HSC-independent lymphopoiesis budding from the ventral endothelium in the zebrafish (*2*) and recently confirmed in the mouse AGM (*5*). Whether these “embryonic multipotent progenitors” correspond to the lymphoid primed nHSPCs identified in this study will require further analysis. Nevertheless, our discovery suggests that their formation is also regulated by the endothelium and specifically by Notch activity in hemogenic cells. This new information could be used to optimize engineering T cells during *ex-vivo* production, for example during CAR-T cells manufacture (*59*).

Reprogramming of somatic cells (including the endothelium) to produce HSPCs with long-term self-renewal and engraftment capacity often lead to cell products with heterogeneous composition, which is not desirable for HSPC transplantation in blood cancer patients (*60-62*). Our discovery suggests that the heterogeneity observed in *ex-vivo* HSPC production might not be an effect of this cost and labor-intensive manufacturing procedure, but an intrinsic property of HSPCs produced by the signaling activated in somatic cells. Indeed, we found that the endothelium of the AGM express inhibitory mechanisms to regulate HSPC heterogeneity, like miR-128. So far, we found that eliminating the miR-128 post-transcriptional regulation of *csnk1a1* and *jag1b*, can differentially control lineage priming bias and cell cycle state dependent HSPC production. Such phenotypes are even more obvious in the AGM of our *csnk1a1* and *jag1b* 3’UTR mutants versus the miR-128 loss alone, suggesting that miR-128 regulates other genetic circuits to further control HSPCs phenotypes. Our work suggests that the rich arsenal of miR-128 target genes can be exploited to dissect and modulate precise HSPC phenotypes in the AGM and to promote the balanced production of HSPCs *ex-vivo*, the holy grail of this life saving application.

## Materiel and methods

### Zebrafish husbandry

Zebrafish were raised and maintained at 28.5C using standard methods, and according to protocols approved by the Yale University Institutional Animal Care and Use Committee (#2017-11473). The following Zebrafish transgenic lines have been described previously: miR-128^ya315-316^ (ZDB-CRISPR-161031-5 and ZDB-CRISPR-161031-9) referred as miR-128^Δ/Δ^, Tg(kdrl:gfp)^zn1^ (ZDB-ALT-070529-1, referred to as kdrl:GFP+), Tg(runx1:eGFP)^y509^ (ZDB-ALT-170717-3, referred to as runx1:GFP), Tg(kdrl:hras-mCherry)^s896^ (ZDB-ALT-081212-4 referred to as kdrl:mCH+); Tg(cmyb:GFP)^zf169^ (ZDB-ALT-071017-1, referred to as cmyb:GFP+), Tg(7xTCF-Xla.Sia:NLS-mCherry)^ia5^ (ZDB-TGCONSTRCT-110113-2 referred to as TCF:NLS-mCh) and (Tg:TP1:eGFP)^um14^ (ZDB-ALT-090625-1 referred to as TP1:eGFP). Zebrafish embryos and adults were genotyped as described (*20*), with primers listed in supplementary table 6.

### Fluorescence activated cell sorting

For all the experiments, 26 hpf embryos were anesthetized with 1X tricaine. Whole, trunk or head dissected embryos were placed in PBS 1X (pH 7.4, Invitrogen) and were dissociated into single cell suspensions through treatment with liberase enzyme (Roche) for 1 hour at 28°C. Liberase was then inactivated with fetal bovine serum (Thermofisher) and cell suspension were washed with cell suspension media (0.5% FBS, 0.8uM CaCl2, 1% Penicillin, leibovitz medium L15 (Gibco)) (*63*). DAPI was added to the single cell suspension to distinguish alive cells. We recovered FAC-sorted cells in pre-coated 1.5ml tubes with 0.04% BSA in PBS.

### Quantitative RT-PCR

Embryos were processed whole or dissociated into single cell suspensions and subjected to FACS. RNA was extracted from ∼300,000 to 600,000 cells using Trizol (Ambion) and 300-500 ng of total RNA was used in a mRNA reverse transcription reactions (Superscript 4, Thermofisher) and the resulting cDNA was used as template for SYBR Green-based quantitative PCR (Kapa biosystems). 100pg-10ng of total RNA was used in miRNA reverse transcription (MirCury LNA miRNA PCR assays, Qiagen), cDNA was used as template for the SYBR Green-based quantitative PCR (MirCury LNA SYBR Green PCR, Qiagen). U6 primers (U6 snRNA), zebrafish miR-128 (dre-miR-128-3p miRCURY LNA miRNA PCR assay) and human miR-128 (hsa-miR-128-3p miRCURY LNA miRNA PCR assay) were used as commercially provided from Qiagen.

The 2^-CT^ method was used to determine relative gene expression for quantitative RT-PCR analyses. mRNA levels were normalized to the beta actin housekeeping gene, *actb1* and was relative to the indicated control, while mature miRNA expression was normalized to U6 snRNA levels and relative to the indicated control. Statistical comparisons between replicate pair CT values for indicated groups were determined by a paired, two-tailed Student’s t-test. All primers are listed in listed in supplementary Table 6.

### Immunofluorescence

Immunofluorescence was performed in all zebrafish stages as follow. After overnight 4% paraformaldehyde (Santa cruz) fixation at 4°C, 27 hpf embryos were washed with 1X PBS-0.01%Tween-20 (PBSTw) 4-6 times for 5 minutes, and then permeabilized with 0.125% trypsin (Millipore Sigma T4549) for 3 minutes. Embryos were washed in blocking solution (0.8% Triton-X, 10% normal goat serum, 1% BSA, 0.01% sodium azide in PBSTw) 3 times for 5 minutes, plus an additional 2-hour incubation with shaking. Antibody concentrations used were 1:300 chicken anti-GFP (Abcam, ab13970, RRID: AB_300798) or rabbit anti-RFP (Antibodies Online, ABIN129578, RRID: AB_10781500) primary antibodies and 1:400 Alexa Fluor 488 goat anti– chicken IgG165 (Thermo Fisher Scientific Cat# A-11039, RRID:AB_2534096) or Alexa Fluor 546 donkey anti-rabbit IgG (Thermo Fisher Scientific Cat# A10040, RRID:AB_2534016) secondary antibodies. Following each overnight antibody incubation at 4°C, six washes for a total of 4 hours were performed with blocking solution lacking goat serum at room temperature and then stained as stated above by immunofluorescence with the addition of DAPI staining (1:500).

### Edu incorporation assay

Click-it EdU Alexa Fluor 647 kit (Thermo Fisher, C10340) was used to analyze S phase endothelial and positively expressing cmyb cells in the 32 hpf AGM or at 4.5 dpf CHT. Embryos were injected at 32 hpf or 4.5 dpf with 10mM Edu staining solution into the sinus venosus and incubated for 5 minutes at 28C, followed by 4% paraformaldehyde overnight fixation at 4C. Embryos were washed 3 times for 5 minutes with PBSTw and placed in cold 100% acetone at −20C for 7 minutes and rinsed with dH_2_O. Subsequently, embryos were permeabilized with 1% DMSO, 1% Triton in 1xPBS for an hour and washed 3 times for 5 minutes with PBSTw and then incubated with reaction cocktail (1x reaction buffer, CuSO_4_ solution, Alexa Fluor azide and reaction buffer additive) for 1 hour at room temperature in the dark. Samples were rinsed 5 times for 5 minutes with PBSTw, and then stained as stated above by immunofluorescence with the addition of DAPI staining (1:500).

### Whole mount in situ hybridization

Whole mount in situ hybridization (WISH) with probes against *gata2b, runx1, cmyb, gata1a, ikaros, lcp1, rag1, notch3, flt4, etv2 and scl* were performed as previously described (*20*). Briefly, Embryos were fixed in paraformaldehyde 4% overnight and washed in methanol. Embryos were kept at −20C. Embryos were then rehydrated with PBTw and permeabilized with 10ug/ml proteinase K (Roche) (10 minutes for 24 hpf, 13 minutes for 32 hpf, 1 hour for 4.5 dpf), followed by a post-fixation in paraformaldehyde 4% for 20 minutes. Then embryos were incubated with the specific probes overnight at 65C. Finally after extensive wash, embryos were incubated with anti-DIG antibody 1:10,000 (Roche). Imaged embryos were quantified as follows: *gata2b, cmyb, runx1* stained cells were counted in the region of the dorsal aorta above the yolk extension; *lcp1* and *gata1a* stained cells were counted in the CHT at 4.5 dpf. *ikaros* and *cmyb* staining at 4.5 dpf and 3 or 6 dpf were quantify as area of staining using ImageJ. Bright-field images of WISH staining were acquired with a Leica Microsystems M165FC stereomicroscope equipped with Leica DFC295 camera.

### Image acquisition and analysis

Zebrafish embryos were treated with 0.003% 1-phenyl-2-thiourea (PTU, Sigma P7629) starting at 70/80% gastrulation stage to prevent pigmentation. Embryos imaged live by confocal microscopy were anesthetized in 0.1% tricaine and mounted in 1% low melt agarose. Fluorescent images and time-lapse movies were captured using Zeiss LSM 980 confocal microscope using 20X water immersion objective. Confocal time-lapse movies were performed at room temperature starting at 27 hpf with z-stacks acquired at an interval of 12 minutes for a total of 15 hours. Time of delamination was quantified for *cmyb:GFP*+ *kdrl:mCherry*+ cells transitioning from flat morphology until they exit the ventral dorsal aorta wall into the subaortic space. Cells were tracked using dragon-tail analysis in Imaris software (V.9.9.1, Bitplane, United Kingdom).

### Flow cytometry of Whole Kidney Marrow (WKM)

Adult WKM (1 month-old) or trunks (26 hpf) were mechanically dissociated (*64*). DAPI was used to differentiate alive cells. Cells were analyzed using a LSR Fortessa (BD Biosciences). Quantification was done through FlowJo software.

### miR-128 expression construct

The miR-128 endothelial expression construct ws constructed as previously described with the following modifications (*20*). To generate the pME-miR-128 middle entry cassette for Gateway-compatible cloning, a 365 bp genomic sequence containing the miR-128-1 stem loop precursor was PCR amplified with flanking KpnI and StuI sites, and the resulting fragment was restriction digested and cloned into pME-miR (p512 addgene) using T4 ligation (NEB M0202S) according to the manufacturer’s protocol. Promoter entry cassette p5E-fli1a (Addgene #31160) were utilized in LR multisite Gateway cloning reactions as performed before (*20*) to produce fli1a:mCherry-miR-128 constructs, respectively. Embryos were injected in the one-cell with 25 pg of the expression construct and Tol2 transposase mRNA, and later selected for mCherry expression.

### gRNA generation & injection

CRISPRScan (https://www.crisprscan.org) was used to design gRNAs to mutate the miR-128 responsive element region in jag1b and csnk1a1 3’UTRs with CRISPR/Cas9 genome editing. gRNA preparation was performed according to published protocols with the following modifications (*65*). F0 embryos were injected with 100 pg of gRNA and 200 pg of Cas9 mRNA at the one-cell stage. T7E1 assay or PCR amplification were used to validate mutations and were analyzed in phenotypic assays. Sequences and primers are listed in supplementary table 6.

### Statistical analysis

All images of embryo embryos were blinded before quantification. Blind analysis ensures that all analytical evaluation were independent from genotype or treatment. All experiments were performed at least with 3 independent experiments or replicas. A two-tailed Mann-Whitney U test was applied for low biological replicates per genotype to detect statistical differences between conditions due to the relatively low sample numbers (less than 8 replicas and 5-20 embryo/cells each replica and condition). A two-tailed Mann-Whitney U test was used also when the dataset was not distributed normally as determined with a Shapiro-Wilk test using the default parameters in GraphPad Prism 7.

### Bulk and single cells RNA sequencing sample preparation

To identify the vascular transcripts regulated by miR-128, 3 replicates of wild-type or miR-128 mutant *kdrl:GFP*^+^ endothelial cells were FAC-sorted from 26 hpf dissected head and trunk tissue as in (*63*). Total RNA was then isolated with the Lexogen SPLIT RNA Extraction Kit, and ∼ 10 ng was used to prepare Lexogen QuantSeq 3′ mRNA-Seq libraries for Illumina deep-sequencing according to the manufacturer’s protocol. Libraries were amplified with ∼ 17 PCR cycles using the Lexogen PCR Add-on Kit according to the Lexogen manufacturer’s protocol.

CD34+CD43-hPSC-derived cells were sorted on day 8 of differentiation and were RNA extracted with Trizol. Libraries (Trunk: 3 for WT and 4 for miR-128^Δ/Δ^, Head: 2 for WT and 4 for miR-128^Δ/Δ^) were then prepared the same way as Zebrafish cells. 5 ng of RNA were used in each sample and 19-21 cycles were used for libraries amplification. Libraries were then deep sequenced according to the Illumina manufacturer’s protocol on an illumine Hiseq 2500, at the Yale Center for Genome Analysis. Wild-type, miR-128^Δ/Δ^, *csnk1a1* and *jag1b* g3’UTR mutants Tg(kdrl:GFP)^zn1^ trunk tissue containing the AGM were dissected at 26 hpf. Dissected trunk tissues were dissociated into single cell suspensions and subjected to FACS. GFP+ cells, which had > 85% cell viability, were loaded onto the 10X Genomics Chromium instrument for a targeted recovery of 10,000 cells per sample. 10X Genomics Chromium Next GEM Single Cell 3’ Library Construction Kit V3.1 (CG000204) was used to generate libraires according to manufacturer instructions. Barcoded libraries were sequenced on an Illumina HiSeq 4000 instrument.

### Whole RNA sequencing analysis

Whole transcriptome profile (bulk RNA sequencing) was analyzed with principal workflow demonstrated by Lexogen. Freely available tools were part of the Galaxy platform (*66*). Specifically, quality of data was checked with FastQC (*67*). BBDuk was used to remove the adaptor contamination, polyA readthrough, and low-quality tails (*68*). Zebrafish genome index was generated with STAR (*69*) according to GRCz10 (Ensembl release 91) (*70*), and decontaminated reads were mapped to the zebrafish genome (*71*). Output BAM files were indexed with SAMtools (*72*). Reads were counted with HTSeq (*73, 74*). Genes below 5 read counts in all replicates in either condition were filtered out with a customized python script. Differentially expressed genes between WT and miR-128-/- mutant conditions were identified with DESeq2 (*75*). Significantly differentially expressed genes in miR-128-/- trunk and head endothelial cells were examined for miR-128 binding sites with TargetScanFish Release 6.2 (*76*). KEGG pathway terms were assigned to differentially expressed genes with DAVID (*77, 78*). Differentially expressed miR-128 target genes, KEGG pathways can be found in supplementary Table 2.

### Human Whole RNA-SEQ Processing

Samples were processed using a similar pipeline as described for zebrafish samples above. No Poly-A sequence removal was necessary. Ensemble genome reference build GRCh38 was used. WNT- and NOTCH-regulated genes are shown in supplementary Table 4.

### Single cell RNA sequencing analysis

#### Single-cell RNA sequencing, quality control

RNA sequencing quality assurance was performed using *FastQC* (*Version 0*.*11*.*9*), by looking for the presence of adapters and sequence quality through Phred Score. Genome alignment was performed using the *10X* Genomics *Cell Ranger* pipeline (*Version 5*.*0*.*0*). A transcriptome reference using a customized zebrafish genome annotation (*79*) was built that corrected 3’ UTR annotation problems and improved alignment performance. The resulting filtered feature-barcode matrices were used for downstream analysis.

The filtered count matrices were loaded on *RStudio* (*Version 4*.*1*.*1*), and the *Seurat* (*Version 4*.*0*.*6*) class object was used to store the data. Cells with less than 200 features and features detected in less than three cells were removed. Cell quality control was performed looking at the overall distribution of counts, detected genes, and expression of mitochondrial genes.

Cell doublets, cells having more than 35,000 Unique Molecular Identifiers (UMI), were removed. After those quality controls, 22,230 cells (WT and miR-128^Δ/Δ^ cells) were kept. Each sample was integrated and batch effect correction was performed using an algorithm based on Mutual-Nearest Neighbors, which finds shared cell populations across different datasets and creates anchors to remove non-biological signals (*80*). For this, data were normalized with *NormalizeData* and 2000 High Variable Genes (HVG) were identified. These HVG were used to define the integration anchors. The first 20 dimensions were used to perform the integration.

#### Single-cell RNA sequencing, Data Analysis

Uniform Manifold Approximation and Projection (UMAP) was applied to the integrated data to obtain a representation of the manifold. The neighborhood graph was calculated using *FindNeighbors* and clusters were extracted using *FindClusters* at the resolution of 0.5, both functions defined on *Seurat*.

Cell type identities were assigned to clusters through visualization of canonical cell type markers and differential gene expression. Gene markers defining each cell cluster regardless of cell genotype were identified using the *FindConservedMarkers* function implemented on *Seurat*. EHT cell clusters, 6,096 cells were then subset and re-clustered. The cell subset was re-analyzed, rescaled, and normalized, new HVG were identified and new UMAP with a resolution of 0.3 was produced.

#### Module analysis

*Monocle 3 find_gene_modules* function was employed to explore cell identities by finding modules of genes co-expressed across cells (*81*). Then, *g:Profiler* was used to functionally characterize the modules through an enrichment analysis. Cell trajectory was also reconstructed by calculating the pseudotime value having as root or starting point the pre-hem C1 population.

#### RNA velocity Analysis

This technique allows to predict the future state of cells through their gene expression based on trends in the unspliced/spliced ratio. *Velocyto* (*Version 0*.*17*.*17*) was used to obtain the unspliced count matrix for each sample (*26*). The count matrices were integrated into the data using the *Anndata* structure on *PyCharm (Version 2021*.*1). scVelo* (*Version* 0.2.3) was used for splicing ratio, where all cells have the same splicing ratio (*82*).

#### Cell cycle Analysis

Cell cycle was characterized by performing a cell cycle scoring analysis using Seurat’s function *CellCycleScoring* with cell cycle phase markers (*83*).

#### Wnt and Notch signaling signature analysis

*AUCell* (*Version 1*.*14*.*0*) was employed to WT and miR-128^Δ/Δ^ cells. The area under the curve (AUC) was used to quantify and test the signature enrichment of Wnt and Notch gene set in each cell. The gene sets used were described in the Kyoto Encyclopedia of Genes and Genomes (*KEGG)* for both Notch (*dre04330*) and Wnt (*dre04310*).

#### Analysis of csnk1a1 and jag1b g3’UTR RNA Samples

The *csnk1a1 and jag1b g3’UTR* single cell samples were processed similarly to the previous ones and resulted in 20,600 cells. Clustering analysis was performed using a resolution of 0.5 and EHT clusters were re-clustered, accounting for a total of 7,949 cells. Cells were projected on the re-clustered UMAP and their identities were predicted using our previous cluster-cell type annotation using *Seurat MapQuery* function. Then, cells were filtered based on their cell type prediction score, removing cells with a *max*.*prediction*.*score* lower than 0.55, followed by a new round of projection. A specific filter (*prediction*.*score* > 0.70) was applied specifically in the cells from clusters 7 and 8 removing cells sparsely distributed throughout the UMAP visualization. In total, 5,782 cells were kept.

#### Sample-associated density estimate and relative likelihood, MELD analysis

MELD was employed (standard parameters) for quantifying the effect of experimental perturbations at the single-cell resolution for a mutant dataset compared to a reference one (*57*). MELD provides a density estimate for each cell in each dataset which allows to identify the cells most or least affected by the perturbation indicating changes in their abundance.

### hPSC differentiation procedure

#### Maintenance and differentiation

The hESC H1 line was maintained on irradiated mouse embryonic fibroblasts in hESC media as described previously (*31, 84*). For differentiation, hPSCs were cultured on Matrigel-coated plasticware (Corning Life Sciences) for 24 hours, followed by embryoid body (EB) generation, as described previously (*30, 62, 85*). Briefly, hPSCs were dissociated with brief trypsin-EDTA (0.05%) treatment, followed by scraping. For the first 3 days of differentiation, EBs were resuspended in SFD media (*86*) supplemented with L-glutamine (2 mM), ascorbic acid (1 mM), monothioglycerol (MTG, 4×10-4 M; Sigma), and transferrin (150 μg/mL). On Day 0, EBs were treated with BMP4 (10 ng/mL). 24 hours later, bFGF (5 ng/mL) was added. On the second day of differentiation, ACTIVIN A (1 ng/mL), SB-431542 (6 μM), CHIR99021 (3 μM), and/or IWP2 (3 μM) were added, as indicated. On the third day, differentiation cultures were changed to StemPro-34 media supplemented with L-glutamine, ascorbic acid, MTG and transferrin, as above, with additional bFGF (5 ng/mL) and VEGF (15 ng/mL). On day 6, IL-6 (10 ng/mL), IGF-1 (25 ng/mL), IL-11 (5 ng/mL), SCF (50 ng/mL), EPO (2 U/mL final). All differentiation cultures were maintained at 37°C. All embryoid bodies and mesodermal aggregates were cultured in a 5% CO2/5% O2/90% N2 environment. All recombinant factors are human and were obtained from Biotechne. Analysis of hematopoietic colony potential via Methocult (H4034; Stem Cell Technologies) was performed as described previously(*30, 32*).

#### miR-128 manipulation within cultures

Cells were left untreated (vehicle control) or treated with antagomiR specific to miR-128 or a scramble antagomir (100 uM in molecular grade water) to a final concentration of 200 nM.

#### Flow Cytometry and Cell Sorting

Cultures were dissociated to single cells, as previously described (*31*). Cells were washed, labeled, sorted and collected in StemPro-34 media. The antibodies used are all as previously described (*30-32*). KDR-PE (clone 89106), CD34-PE-Cy7 (clone 8G12), CD43-FITC (clone 1G10), CD73-PE (clone AD2), CXCR4-APC (clone 12G5), and CD235a-APC (clone HIR-2). All antibodies were obtained from BD Biosciences except for KDR (Biotechne). Cells were sorted with a FACSMelody (BD) cell sorter. For isolation of mesodermal populations, day 3 of differentiation WNTd KDR+CD235a– or WNTi KDR+CD235a+ were FACS-isolated and reaggregated at 250,000 cells/mL in day 3 media, as above. Cultures were plated in 250 μL volumes in a 24 well low-adherence culture plate, and grown overnight in a 37°C incubator, with a 5% CO2/5% O2/90% N2 environment. On day 4, an additional 1 mL of antagomiR- or scramble-supplemented day 3 media was added to reaggregates. On day 6 of differentiation, cultures were fed as normally and subsequent CD34+CD43-cells were sorted on day 8 of differentiation.

#### Hemato-endothelial growth conditions of hPSC-derived hemogenic endothelium

Either CD34+CD43– cells (stage 1 antagomir manipulation) or CD34+CD43-CD73-CD184-cells (stage 2 antagomir manipulation) were isolated by FACS and allowed to undergo the endothelial-to-hematopoietic transition (EHT) as described previously (*32, 62*). Briefly, cells were aggregated overnight at a density of 2×105 cells/mL in StemPro-34 media supplemented with L-glutamine (2 mM), ascorbic acid (1 mM), monothioglycerol (MTG, 4 × 10−4 M; Sigma-Aldrich), holo-transferrin (150 μg/mL), TPO (30 ng/mL), IL-3 (30 ng/mL), SCF (100 ng/mL), IL-6 (10 ng/mL), IL-11 (5 ng/mL), IGF-1 (25 ng/mL), EPO (2 U/mL), VEGF (5 ng/mL), bFGF (5 ng/mL), BMP4 (10 ng/mL), FLT3L (10 ng/mL), and SHH (20 ng/mL). Aggregates were spotted onto Matrigel-coated plasticware and were cultured for additional 9 days. Cultures were maintained in a 37°C incubator, in a 5% CO2/5% O2/90% N2 environment. All resultant cells within the hemato-endothelial cultures were subsequently harvested by trypsinization, and assessed for hematopoietic potential by Methocult in a 37°C incubator, in a 5% CO2/air environment.

## Supporting information

Supplementary Movie 1.A. Time lapse imaging movies from 24 to 50 hpf of WT. Tg(kdrl:mCherrys896,cmyb:GFPzf169).

Supplementary Movie 1.B. Time lapse imaging movies from 24 to 50 hpf of miR-128-/-. Tg(kdrl:mCherrys896,cmyb:GFPzf169).

Supplementary Table 1. Cell numbers and gene expression from single cell RNA sequencing experiment between wild-type and miR-128-/-.

Supplementary Table 2. Raw gene expression and gene ontology on differential gene between wild-type and miR-128-/- from bulk RNA sequencing.

Supplementary Table 3. Total cell numbers and gene expression from single cell RNA sequencing experiment between wild-type, miR-128-/-.

Supplementary Table 4. Raw gene expression of wnt- and notch-regulated genes from bulk RNA sequencing between scramble and antagomir-128 treated hPSCs

Supplementary Table 6. List of primers used in this study.

## Data Availability & Reproducibility

The visualizations were generated using *ggplot2* (V*ersion 3*.*3*.*5*) and *Prism (Version 9)*. The raw fastq files will be available on GEO upon publication. The expression matrices will be available on the genome browser upon publication.

## Acknowledgments

We thank you all the members of the Nicoli Lab for the critical discussion. We also thank you Drs. Dionna Kasper (Geisel School of Medicine at Dartmouth, Molecular and Systems Biology) Karen Hirschi (University of Virginia, Cell Biology), and Kaelyn Sumigray (Yale School of Medicine, Genetics) for the critical comments and discussion of the manuscript.

## Figure Legends

**Supplementary Figure. 1.**
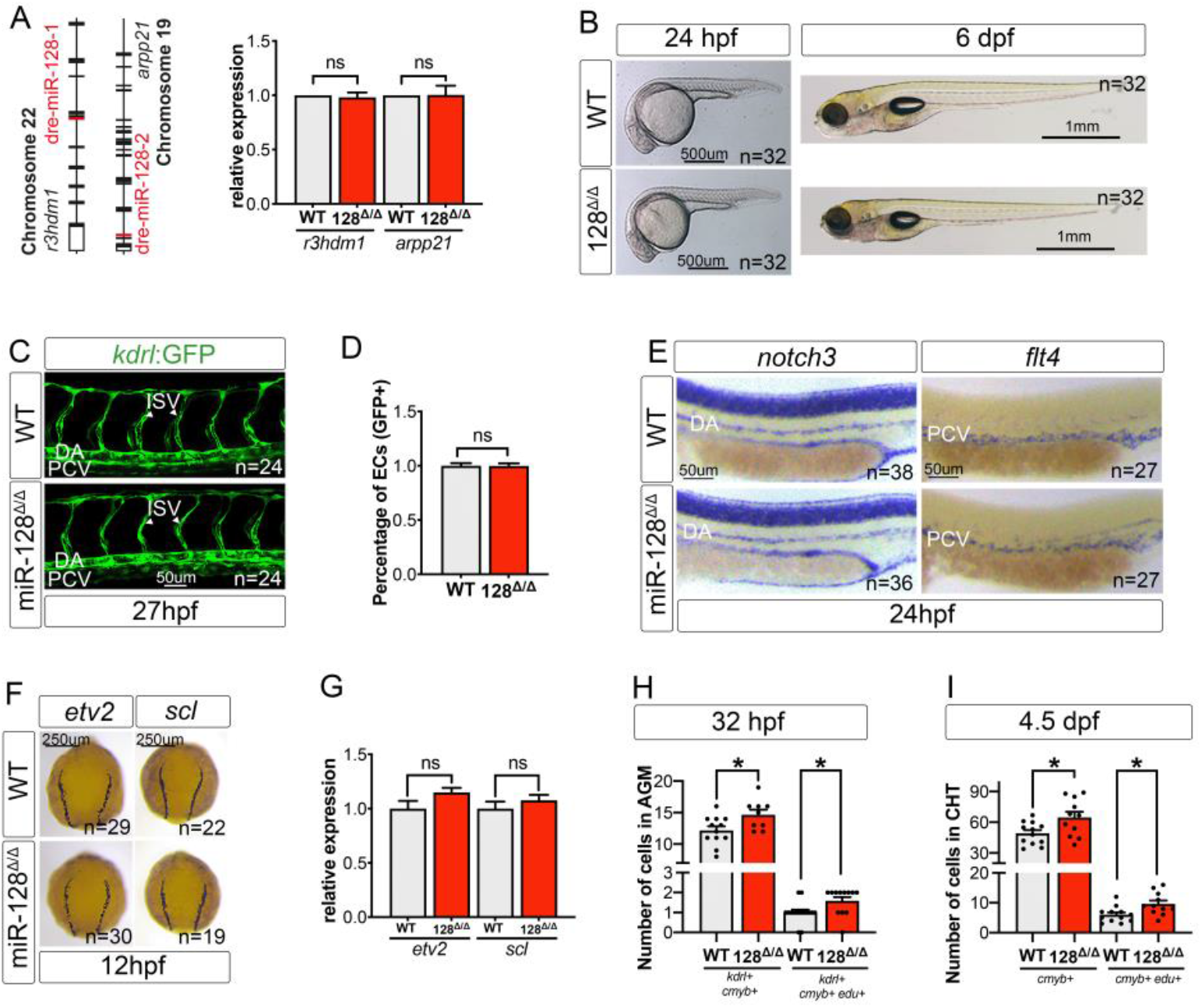
miR-128 does not show any morphological or vascular phenotype while regulating EHT. (A) Schematic representation of the genomic location of *miR-128-1* and *miR-128-2* within *r3hdm1* and *arpp21* introns respectively and qRT-PCR against *r3hdm1* and *arpp21* host genes in WT and miR-128^Δ/Δ^ embryos. (B) Bright-field images of WT and miR-128^Δ/Δ^ embryos at 24 hpf and 6 dpf. (C) Confocal live imaging of Tg(*kdrl:GFP)*^*zn1*^ WT and miR-128^Δ/Δ^ embryos at 27 hpf, showing no vascular defects. (D) Percentage of endothelial cells GFP *kdrl*+ over the total number of trunk cells in WT and miR-128^Δ/Δ^ at 26 hpf assessed by flow cytometry. (E) Arterial and Venus specification are not affected in miR-128^Δ/Δ^ as found by WISH against *notch3* and *flt4* at 24 hpf respectively. (F) bright field images of WISH of the early hematopoietic and endothelial development markers *etv2* and *scl* at 12 hpf (dorsal view). (G) qRT-PCR against *etv2* and *scl* at 12 hpf in miR-128^Δ/Δ^ and WT. (H,I) Number of nHSPCs in miR-128^Δ/Δ^ and WT, assessed by immuno-fluorescence at 32 hpf (H, *kdrl*^+^,*cmyb*^+^ and *kdrl*^+^,*cmyb*^+^,Edu^+^) in the AGM and 4.5 dpf (I, *cmyb*^+^ *and cmyb*^+^,Edu^+^) in the CHT. All quantifications are represented with mean ± SEM. ns: p>0.05, *p≤0.05. Abbreviations: aorta gonad mesonephros (AGM), dorsal aorta (DA), posterior cardinal vein (PCV), caudal hematopoietic tissue (CHT) and intra segmental vessel (ISV).

**Supplementary Figure. 2.**
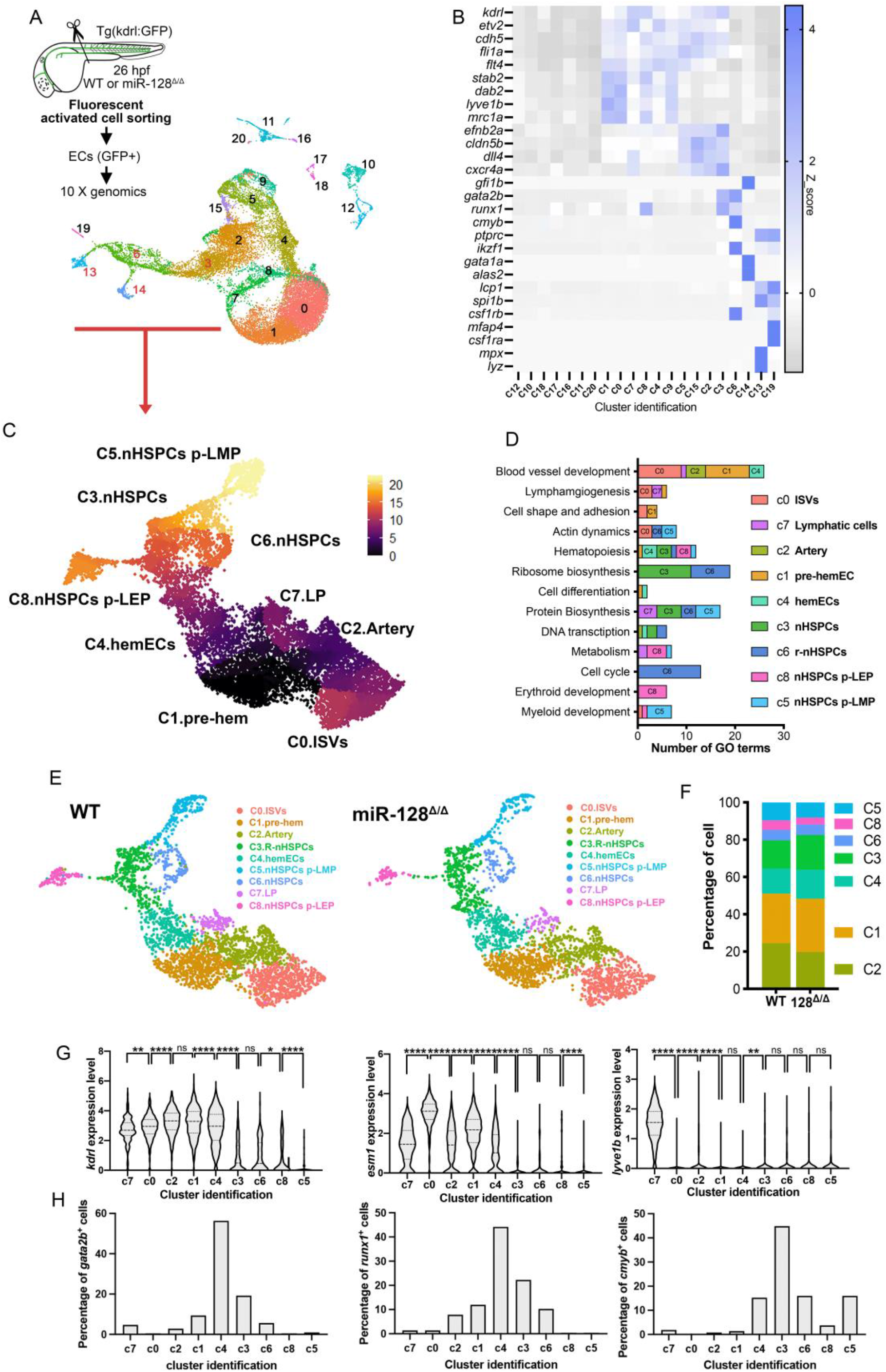
Single cell RNA sequencing identified miR-128 loss of function defects in EHT cell clusters. (A) Schematic representation of single cell RNA-sequencing strategy and UMAP of *kdrl*+ trunk endothelial cells of WT and miR-128^Δ/Δ^ at 26 hpf isolated by FACS from *Tg(kdrl:GFP)*^*zn1*^ line. Cluster numbers in red correspond to the one used for the re-clustering EHT cells in C. (B) Heatmap showing the average gene expression of canonical vascular, hemogenic, blood stem cells and lineage gene markers in single kdrl+ cells. (C) Pseudotime analysis of EHT-clusters for both WT and miR-128^Δ/Δ^ cells (see methods). (D) GO analysis enrichment of EHT-clusters. (E) UMAP of WT and miR-128^Δ/Δ^ of EHT cluster cells with assigned identification. (F) Percentage of cells in each cluster per genotype for the indicated clusters. (G) Violin plot of *lyve1b, esm1* and *kdrl* expression in each cluster of WT cells. (H) Percentage of cells expressing *gata2b, runx1* and *cmyb* in each cluster of WT cells. All Quantification are represented with mean ± SEM. ns: p>0.05, *p≤0.05, **p≤0.01, *** p≤0.001, **** p≤0.0001. Abbreviations: gene ontology (GO).

**Supplementary Fig. 3.**
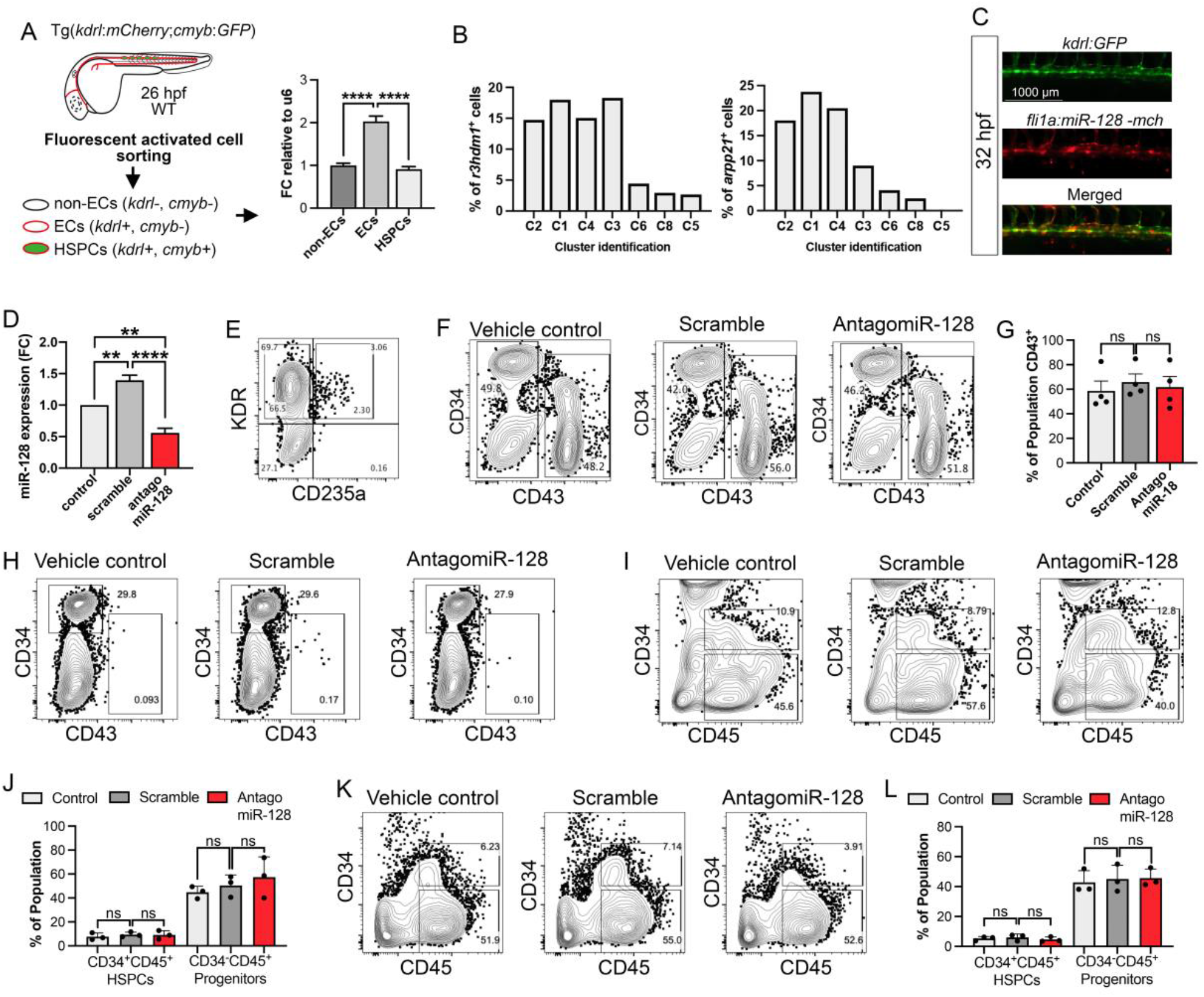
Endothelial miR-128 regulates HSPC heterogeneity. (A) Schematic representation of our HSPCs, endothelia cells (ECs) and non-ECs isolation strategy to quantify miR-128 expression. We used WT *Tg(kdrl:mCherry*^*s896*^*;cmyb:GFP*^*zf169*^) where we separated the non-ECs, ECs and HSPCs at 26 hpf. RT-qPCR against miR-128 and U6 in the FAC-sorted cell showing mature miR-128 enrichment in ECs vs HSPCs or non-ECs. (B) Percentage of cells in scRNA-seq clusters expressing *r3hdm1* and *arpp21* in which the intronic miR-128 gene is hosted. (C) Confocal representative images of *fli1a:miR-128-mCherry* at 32hpf in WT Tg(*kdrl:GFP*^*zn1*^) embryos. mCherry indicates successful miR-128 expression in GFP *kdrl*+ cells in the trunk of 32 hpf zebrafish. (D) Mature miR-128 qRT-PCR expression in hemogenic endothelial cells from human pluripotent stem cell (hPSC) treated with scramble or antagomiR-128. (E) Representative flow cytometry analysis of KDR and CD235a expression during differentiation. (F) Representative flow cytometry analysis of CD34 and CD43 expression during differentiation of cells treated with vehicle control or scramble or antagomiR-128. (G) Quantification of CD43^+^ cells (percentage). (H) Representative flow cytometry analysis of CD34 and CD43 expression during differentiation of cells treated with vehicle control or scramble or antagomir-128. (I) Representative flow cytometry analysis of CD34 and CD45 expression during differentiation of cells treated with vehicle control or scramble or antagomir-128. (J) Quantification of CD34^+^CD45^+^ (HSPCs) and CD34^−^CD45^+^ (progenitors) in percentage. (K) Representative flow cytometry analysis of CD34 and CD45 expression during differentiation of cells treated with vehicle control or scramble or antagomiR-128. (L) Quantification of CD34^+^CD45^+^ (HSPCs) and CD34^−^CD45^+^ (progenitors) in percentage. All quantification are represented with mean ± SEM. **p≤0.01, **** p≤0.0001. Abbreviations: hemogenic endothelium (HE).

**Supplementary Fig. 4.**
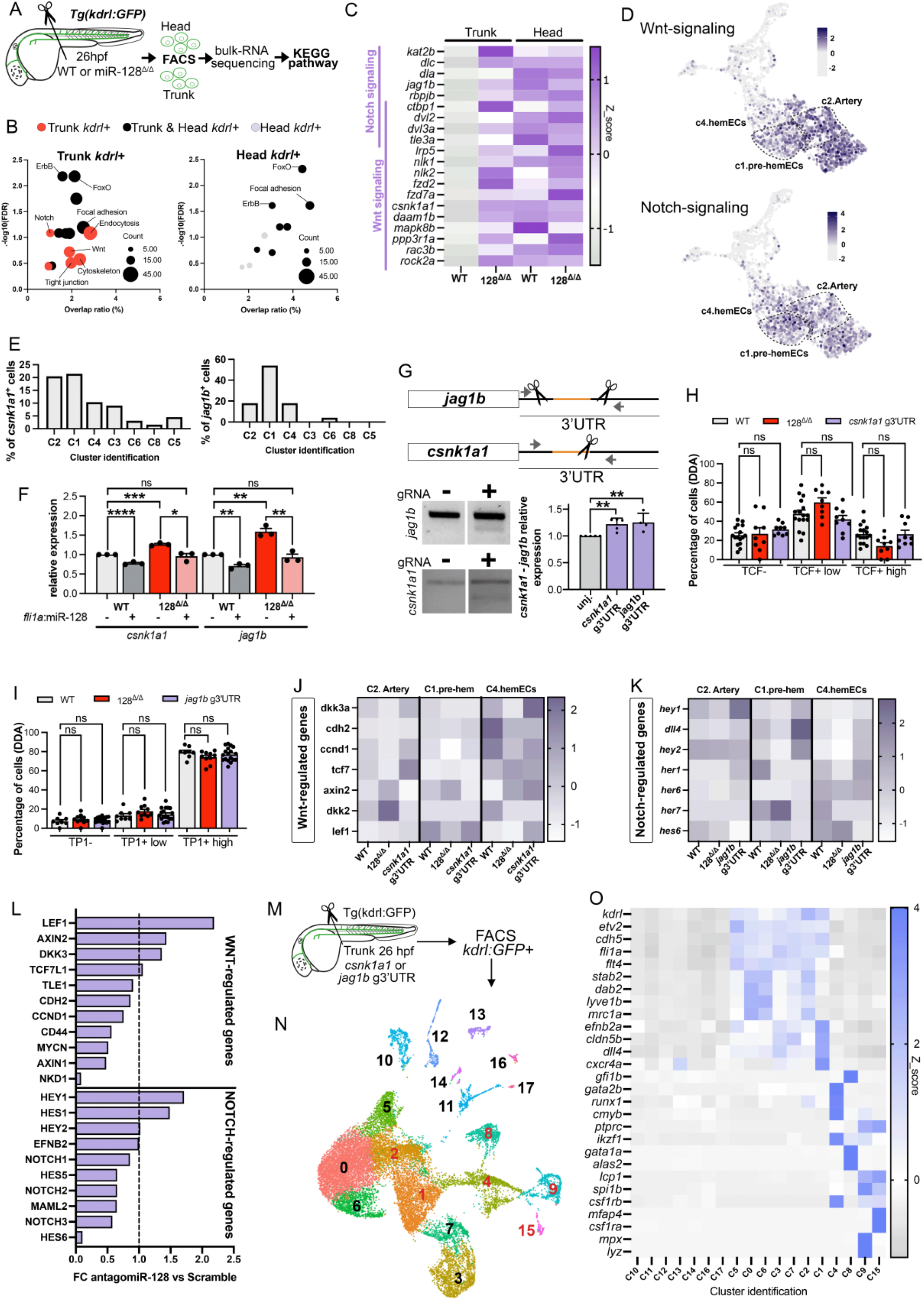
miR-128 controls HSPC heterogeneity by regulating Wnt-csnk1a1 and Notch-jag1b outputs prior EHT. (A) Representation of bulk RNA sequencing on *kdrl*+ endothelial cells of head and trunk at 26 hpf of WT and miR-128^Δ/Δ^ Tg(*kdrl:*GFP^*zn1*^) embryos. (B) KEGG pathway analysis of differentially expressed miR-128 target genes in trunk or head *kdrl*+ cells. Wnt and Notch signaling are specific to miR-128^Δ/Δ^ trunk *kdrl*+ cells. (C) Heatmap of bulk RNA sequencing in endothelial cells of trunk or head of WT and miR-128^Δ/Δ^ putative target genes within the Notch and canonical Wnt signaling at 26 hpf. (D) UMAP of Wnt- and Notch-signaling signature in wild-type cells (see methods). (E) Number of cells expressing *csnk1a1* or *jag1b* in scRNA-seq WT EHT clusters. Percentages represent the number of cells expressing *csnk1a1* or *jag1b* within the total number of cells. (F) qRT-PCR validation of miR128-*csnk1a1* and -*jag1b* regulation. Both expressions were increased in miR-128^Δ/Δ^ and rescued by *fli1a:miR-128* (+). (G) Schematic of *csnk1a1* and *jag1b* Cas9 and guide RNA design on the genomic locus of the gene respective 3UTR. PCR and T7 endonuclease test was used to validate generation of indels after 24 h from Cas9 and *csnk1a1* and *jag1b* gRNA injection respectively. De-repression of *csnk1a1* and *jag1b* were validated by qRT-PCR at 24 hpf. (H-I) Quantification of TCF-mCherry (H) and TP1-GFP (I) responsive cells in the dorsal floor of the dorsal aorta (DDA) of WT, miR-128^Δ/Δ^, *csnk1a1* and *jag1b* g3’UTR. (J) Heatmaps of canonical Wnt-responsive genes normalized with z-score in WT, miR-128^Δ/Δ^ and *csnk1a1* g3’UTR per clusters. (K) Heatmaps of Notch-responsive genes normalized with z-score in WT, miR-128^Δ/Δ^ and *jag1b* g3’UTR per clusters. (L) Bar graph representing average fold change expression of WNT- and NOTCH-regulated genes assessed by whole-transcriptome RNA-sequencing of hPSC-derived WNTd CD34+CD43-cells following scramble or antagomiR-128 treatment during stage 1. (M) Schematic representation of single cell RNA sequencing strategy used in *csnk1a1* and *jag1b* g3’UTR and UMAP of trunk *kdrl*+ endothelial cells at 26 hpf. (N) UMAP of *csnk1a1* and *jag1b* g3’UTR cells. Clusters indicated with red numbers correspond to EHT cells re-clustered in the main figure. (O) Heatmap of vascular, hemogenic, blood stem cells and blood lineage cell defining markers. All quantification are represented with mean ± SEM. ns: p>0.05, *p≤0.05, **p≤0.01, *** p≤0.001, **** p≤0.0001. Abbreviations: aorta gonad mesonephros (AGM), un-injected (unj.), T7 endonuclease (T7EI).

**Supplementary Table 1**. *Cell numbers and gene expression from single cell RNA sequencing experiment between wild-type and miR-128*^*Δ/Δ*^.

(A) Average expression of markers in all cells within each clusters. (B) Gene ontology terms analysis, obtained by DAVID bioinformatics resource, of cluster defining genes. (C) Total number of cells per clusters in WT and miR-128^Δ/Δ^ cells. Number of cells expressing *gata2b, runx1, cmyb, r3hdm1* and *arpp21* per clusters in WT cells. (D) Raw gene expression in WT and miR-128^Δ/Δ^ cells. (E) Gene ontology terms analysis in each modules defining clusters C3, C6, C8 and C5. (F) Numbers of cells defined by their cell cycle status (G1, G2/M and S phases) in each clusters in WT cells. (G) Number of cells expressing *gata2b*^*+*^ cells in WT and miR-128^Δ/Δ^ cells.

**Supplementary Table 2**. *Raw gene expression and gene ontology on differential gene between wild-type and miR-128*^*Δ/Δ*^ *from bulk RNA sequencing*.

(A) Gene ontologoy terms, obtained by DAVID bioinformatics resource, from differential gene expression between WT and miR-128^Δ/Δ^ trunk endothelial (*kdrl*:*GFP*^+^) cells. (B) Gene ontologoy terms, obtained by DAVID bioinformatics resource, from differential gene expression between WT and miR-128^Δ/Δ^ head endothelial (*kdrl*:*GFP*^+^) cells. (C) Raw gene expression of miR-128 predicted target genes part of Wnt and Notch signaling, in WT and miR-128^Δ/Δ^ endothelial cells from trunk or head bulk RNA sequencing.

**Supplementary Table 3**. *Total cell numbers and gene expression from single cell RNA sequencing experiment between wild-type, miR-128*^*Δ/Δ*^.

(A) Number of cells expressing *csnk1a1*^*+*^ and *jag1b*^*+*^cells in WT cells. (B) Average expression of Wnt- and Notch-regulated genes in each cluster per genotype (WT, miR-128^Δ/Δ^ csnk1a1 and jag1b 3’UTR mutants).

**Supplementary Table 4**. *Raw gene expression of wnt- and notch-regulated genes from bulk RNA sequencing between scramble and antagomir-128 treated human pluripotent stem cells*.

Expression of Wnt- and Notch-regulated genes in scramble or antagomiR-128 treated hPSCs.

**Supplementary Table 5**. *Total cell numbers and gene expression from single cell RNA sequencing experiment between wild-type, csnk1a1 and jag1b 3’UTR mutants*.

(A) Average gene expression in *csnk1a1*- and *jag1b*-3’UTR mutants cells per cluster. (B) Total number of cells in each cluster per genotype (WT, miR-128^Δ/Δ^ *csnk1a1* and *jag1b* 3’UTR mutants). (C) Number of cells expressing *gata2b*^*+*^ cells in WT, miR-128^Δ/Δ^, *csnk1a1* and *jag1b* 3’UTR mutants cells.

**Supplementary Table 6**. *List of primers used in this study*.

**Supplementary Movie 1**. *Time lapse imaging movies from 24 to 50 hpf of WT and miR-128*^*Δ/Δ*^ *Tg(kdrl:mCherry*^*s896*^,*cmyb:GFP*^*zf169*^).

(A) Time-lapse imaging of the AGM region of WT embryo. (B) Time-lapse imaging of the AGM region of miR-128^Δ/Δ^ embryo.

## Notes

### Competing Interest Statement

The authors have declared no competing interest.

