## Supplementary figures and images for "Hematopoietic stem and progenitor cell heterogeneity is inherited from the embryonic hemogenic endothelium"

### Supplementary Movie 1.B. Time lapse imaging movies from 24 to 50 hpf of miR-128-/-. Tg(kdrl:mCherrys896,cmyb:GFPzf169).

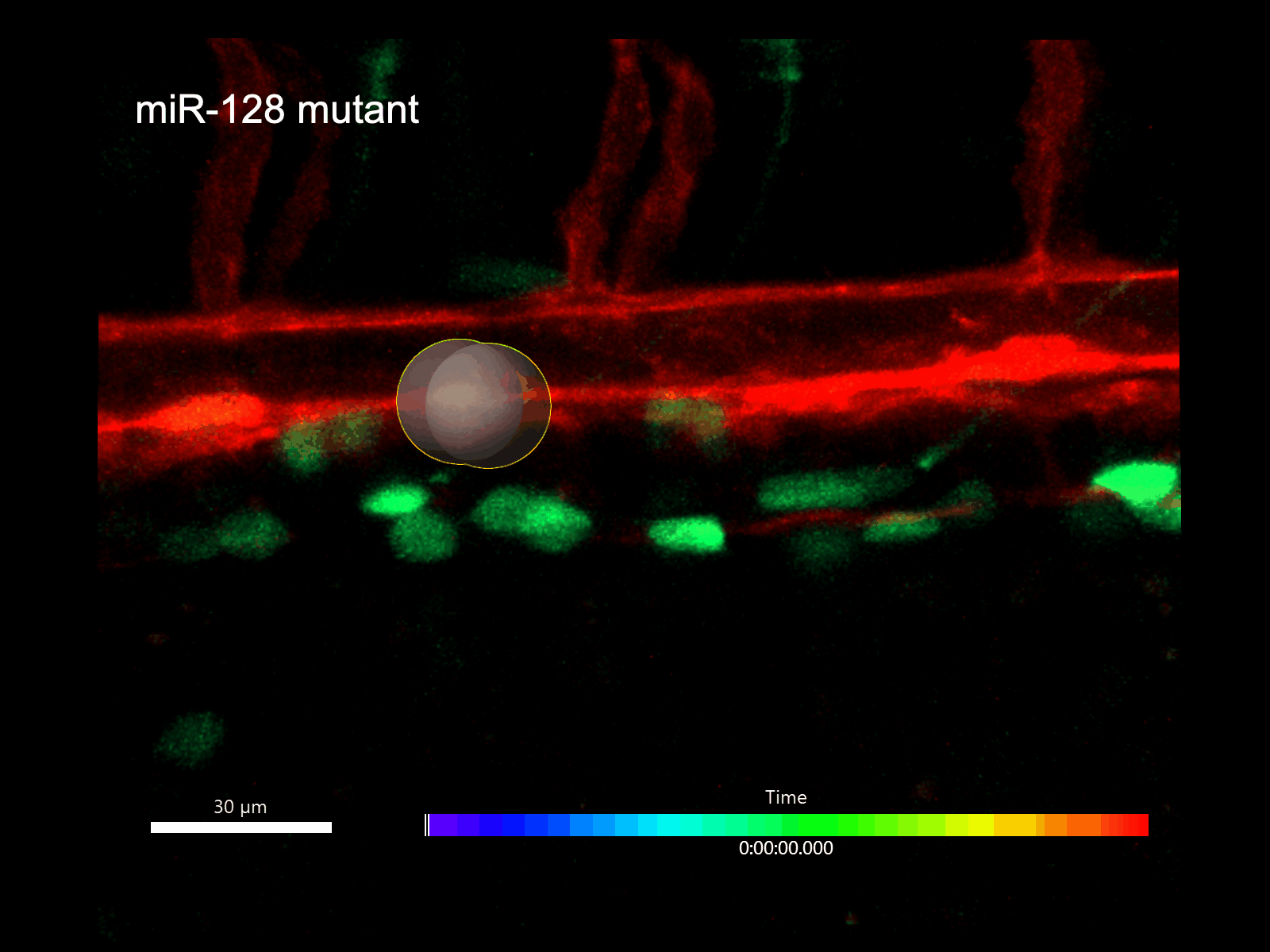
